# Comparative metagenomic analysis provides novel insights into the ecophysiology of Myxococcota in activated sludge systems

**DOI:** 10.1101/2024.04.26.591422

**Authors:** Hazuki Kurashita, Masashi Hatamoto, Shun Tomita, Takashi Yamaguchi, Takashi Narihiro, Kyohei Kuroda

## Abstract

Myxobacteria, belonging to the phylum Myxococcota, are ubiquitous in soil, marine, and other environments. Recent metagenomic sequencing analysis has shown that Myxococcota are predominant in activated sludge systems; however, their ecophysiological traits remain unclear. In this study, we evaluated the potential biological functions of 46 metagenomic bins of Myxococcota reconstructed from activated sludge samples from four municipal sewage treatment plants. The results showed that most Myxococcota bins had an almost complete set of genes associated with glycolysis and the TCA cycle. Palsa-1104 and Polyangiales bins contain glycoside hydrolase GH13 and peptidase M23, which are presumably involved in the lysis of the cell wall and cellular cytoplasm, indicating that part of the Myxococcota from activated sludge can prey on other microorganisms. The previously known cell contact-dependent predatory functions of *Myxococcus xanthus* are conserved in the family Myxococcaceae, though not in other families. Two bins belonging to Palsa-1104 had phototrophic gene clusters, suggesting that these bins may have both heterotrophic and autotrophic metabolisms. Assessment of the social behavior of the activated sludge Myxococcota, a FruA gene, and a C-signal gene involved in the regulation of fruiting body formation were lacking in Myxococcota bins, suggesting that they are incapable of fruiting body formation. In addition, multiple bins of Myxococcota had novel secondary metabolite biosynthesis gene clusters that may be used for the predation of other bacteria in the activated sludge. Our metagenome-based analyses provide novel insights into the microbial interactions associated with Myxococcota in activated sludge ecosystems.

## 1. Introduction

Myxobacteria, which are ubiquitous in soil and oceans, were first described in 1892 (Thaxter, 1892). Myxobacteria belong to the order Myxococcales of the class Deltaproteobacteria; however, recent reclassification of the phylum Proteobacteria has led to the proposal of the phyla Myxococcota, Desulfobacterota, and SAR324 (Langwig et al., 2022; Waite et al., 2020). *Myxococcus xanthus*, the most studied myxobacteria, has a unique life cycle in which myxobacterial cells aggregate and form fruiting bodies upon sensing starvation (referred as social behavior) and produces spores that are resistant to desiccation (Gokhale, 1947; Koch and White, 1998). Additionally, it has a large genome size (up to 15 Mb) (Han et al., 2013; Schneiker et al., 2007) compared to other bacteria and various biosynthetic gene clusters (BGCs) for secondary metabolites such as myxovirescin and myxalamid (Weissman and Müller, 2010).

The formation of fruiting bodies has been confirmed in *M. xanthus*, *Minicystis rosea*, and *Stigmatella aurantiaca* (Garcia et al., 2014; Silakowski et al., 1996). Thirteen modules (enhancer-binding protein (EBP), MrpC, Nla24, exopolysaccharide (EPS) production, FruA, C-signal, A-signal, aggregation-sporulation-fruiting body formation, development timer, adventurous motility, social motility, outer membrane exchange (OME), and chemosensory pathways/rippling) have been well studied as they are the gene modules involved in the social behavior of *M. xanthus* (Murphy et al., 2021). Contrarily, metagenome-assembled genomes of Myxococcota (such as, JAFGVO01, JAFGXQ01, and JAFGIB01) obtained from anaerobic sediments of Zodletone hot springs have been reported to lack genes belonging to the FruA module, C-signal, and EPS production, which are similar to the genera *Anaeromyxobactoer* and *Labilithrix* (Murphy et al., 2021; Thomas et al., 2008; Yamamoto et al., 2014). Zodletone Myxococcota and Anaeromyxobacter lack homologs of sporulation genes (Murphy et al., 2021) and the phenotypic characteristics of *Labilithrix luteola* and *Vulgatibacter incomptus* show no aggregates, fruiting bodies, or spore formation in pure culture (Yamamoto et al., 2014). Therefore, the life cycles of the phylum Myxococcota vary according to their phylogeny and ecophysiology.

Myxococcota are bacterial predators cultivated using macromolecular substrates such as *Escherichia coli* and cellulose (Han et al., 2013; Livingstone et al., 2017). Genomic analysis indicates that it encodes multiple peptidases that degrade cell walls and carbohydrate-active enzymes (CAZymes) (Thiery and Kaimer, 2020). In addition, recent studies have elucidated the cell-contact predatory function of *M. xanthus*, confirming the importance of the tight adherence (Tad) and Type III secretion systems (T3SS) for both the induction of cell death and lysis (C. Wang et al., 2023). The isolates and metagenome-assembled genomes of Myxococcota from anaerobic environments have been reported to contain genes relevant to anaerobic ATP production (such as nitrogen fixation, HydABC, and the Wood-Ljungdahl pathway) and sulfur metabolism (such as QmoABC, AprAB, and DsrAB), which are not present in known aerobic myxobacterial genomes, suggesting that Myxococcota have acquired several functional genes for adaptation to different environments (Murphy et al., 2021). As the metabolic functions of Myxococcota in different environments are diverse, further accumulation of physiological and genomic information through cultivation and genomic analyses is required.

Recent large-scale shotgun metagenomic analyses of activated sludge systems have suggested that Myxococcota with large genome sizes are predominant and have the potential to produce various bioactive substances (Liu et al., 2021). In comparison with other bacteria, Myxococcota has many BGCs, indicating that Myxococcota in activated sludge systems can be useful biological resources, such as biological pesticides. In addition, 16S rRNA-based stable isotope probing (SIP) analyses have revealed the active predatory capabilities of the phyla Myxococcota and Bdellovibrionota in activated sludge systems (Zhang et al., 2023). In particular, Haliangium and mle1-27 (which belong to Myxococcota) exhibit high predatory activity in activated sludge (Zhang et al., 2023). Our previous metagenomic study of 600 activated sludges has also suggested that predatory Myxococcota and Bdellovibrionota contribute to the maintenance of good water quality, such as total carbon and nitrogen removal (Kuroda et al., 2023). However, the detailed ecophysiological functions of Myxococcota remain unclear. Therefore, in this study, we analyzed 46 metagenomic bins of the phylum Myxococcota recovered from activated sludge samples of four wastewater treatment plants (WWTPs E1, F1, G1, and H1) to elucidate the adapted functions of myxobacteria in activated sludge systems and attempted to reveal their ecological roles in activated sludge systems and the potential for secondary metabolite production in the circular bioeconomy.

## 2. Materials and Methods

### 2.1 Sampling, DNA extraction, PCR amplification, and 16S rRNA gene sequence analysis

Activated sludge samples used to treat municipal wastewater were collected from four WWTPs (E1, F1, G1, and H1) in Japan and DNA was extracted using a Fast DNA Spin Kit for Soil (MP Biomedicals, Santa Ana, California, USA) according to the manufacturer’s protocol. The DNA concentration was measured using the Qubit dsDNA BR Assay Kit (QIAGEN). PCR amplification of the 16S rRNA gene was performed using the universal forward primer (Univ515F: 5’-GTGCCAGCMGCCGCGGGTAA-3’) and the reverse primer (Univ909R: 5’-CCCCGYCAATTCMTTTRAGT-3’) (Kozich et al., 2013) for F1, G1, and H1 and (Univ806R: 5’-GGACTACHVGGGTHTCTAAT-3’) for E1 (Caporaso et al., 2012). The PCR products were purified using a QIAquick PCR Purification Kit (Qiagen) according to the manufacturer’s protocol. Purified 16S rRNA gene sequences were analyzed using a MiSeq Reagent Kit v3 and MiSeq system (Illumina, San Diego, CA, USA). Raw 16S rRNA gene sequences were analyzed using QIIME 2 (ver.2021.4) according to previous studies (Kuroda et al., 2023). The 16S rRNA gene sequences were clustered by ≥ 97% similarity to operational taxonomic units (OTUs) using the vsearch software R (Rognes et al., 2016). Taxonomic assignment was performed using classify-sklearn on the SILVA database version 138.

### 2.2 Shotgun metagenomic sequence analysis

Shotgun metagenomic sequencing of the DNA extracted from the activated sludge samples at four WWTPs (E1, F1, G1, and H1) was performed using a NovaSeq6000 (150 bp ×2). Metagenomic analyses of F1, G1, and H1 were performed as described previously (Kuroda et al., 2022). Briefly, raw sequence data were quality-trimmed using Trimmomatic 0.39 (Bolger et al., 2014) and assembled using Megahit2 (Li et al., 2015). Metagenome-assembled bins were generated using the Das Tool 1.1.2 (Sieber et al., 2018) and binning software from Metabat2 (Kang et al., 2019), MaxBin2 (Wu et al., 2016), and MyCC (MyCC_2017. ova) (Lin et al., 2016). The reconstructed bins were assessed for completeness and contamination by using CheckM 1.0.7 (Parks et al., 2015). In this study, we selected bins (belonging to phylum Myxococcota) with ≥80% completeness and ≤10% contamination as high-quality genomes. GTDB-tk ver. 1.5.1 software (r202) was used for taxonomic classification of the bins (Chaumeil et al., 2020).

A genome tree was constructed using the concatenated phylogenetic marker genes of the obtained bin and the phylum Myxococcota. Conserved maker genes were identified using “gtdbtk identify” with default parameters and aligned to reference genomes using “gtdbtk align” with taxonomic filters (-taxa_filter p Myxococcota, p Myxococcota_A, p Myxococcota_B) (Chaumeil et al., 2020). A phylogenetic tree was constructed using IQ-TREE version 2.1.4-beta (-B 1000) with an automatically optimized substitution model (LG+F+R10) (Minh et al., 2020).

### 2.3 Estimation of secondary metabolite synthetic gene in Myxobacteria from activated sludge

All bins belonging to phylum Myxococcota were annotated through a combination of Prokka v1.14.6 (Seemann, 2014), DRAM software (Shaffer et al., 2020), and GhostKOALA (Kanehisa et al., 2016) and manual annotation. To predict secondary metabolite gene clusters, we used antiSMASH (version 5.0 (Blin et al., 2019). In addition, Natural Product Domain Seeker (NaPDoS) (Ziemert et al., 2012) was used to identify the C domains in non-ribosomal peptide synthase (NRPS) and the KS domains in polyketide synthase (PKS) of Myxococcota bins with default parameters. The C and KS domains were subjected to phylogenetic analysis using the NaPDoS to generate phylogenetic trees. To correctly estimate the phylogenetic positions of the KS domain sequences, we excluded genes involved in fatty acid synthesis and polyunsaturated fatty acid synthesis. To assess the social behavior and cell-contact-dependent prey killing of Myxococcota bins obtained from the activated sludge samples, a BLASTP-based homology search (e-value < 1e-5, qcovs > 50%) was performed against the *M. xanthus* DK1622 genome (GCA_000012685.1). Principal component analyses (PCA) based on glycoside hydrolases (GHs) and peptidases were performed using PAST4 software (Hammer et al., 2001). Bins with Calvin–Benson–Bassham (CBB) cycles were selected and annotated with photosynthetic gene clusters using the default parameters from the eggNOG (Cantalapiedra et al., 2021) and KEGG software. Genes involved in phosphate accumulation in polyphosphate-accumulating bacteria were referenced, and the presence or absence of polyphosphate accumulation was assessed using a BLASTP-based homology search (e-value<1e^-5^, qcovs > 50%) and GhostKOALA.

### 2.4 Deposition of DNA sequence data

Raw sequence data from activated sludge samples F1, G1, and H1 have been deposited in the DDBJ Sequence Read Archive (SRA) database (DRA018446). Sequence data from activated sludge sample E1 were obtained from the DDBJ SRA database (DRA015582 and DRA016086) deposited in a previous study (Kuroda et al., 2023). Myxococcota bins data from this study are available at figshare: https://doi.org/10.6084/m9.figshare.25702542.

## 3. Results and Discussion

### 3.1 Detection of Myxococcota recovered from activated sludge systems through 16S rRNA gene and metagenomic sequence analyses

Microbiome analyses based on the 16S rRNA gene and shotgun metagenomic sequencing were performed to assess the ecology of myxobacteria in the activated sludge systems from four wastewater treatment plants (E1, F1, G1, and H1). The phylum Myxococcota was present in the activated sludge at abundances ranging from 3.6 to 9.5% of the total prokaryotes (**Table S1**). Through metagenomic sequencing, 46 metagenomic bins belonging to the phylum Myxococcota (average completeness: 90.5%; average contamination: 5.06%) of the 340 bins (average completeness: 91.6%; average contamination: 3.40%) were recovered. These 46 bins were classified to orders Polyangiales (31 bins), Palsa-1104 (8 bins), Haliangiales (2 bins), Nannocystales (1 bin), Myxococcales (1 bin), UBA796 (1 bin), PHBI796 (1 bin), PHBI01 (1 bin), and UBA9042 (1 bin) based on GTDB-based phylogenies (Chaumeil et al., 2020) (**Table 1, Table S2, Figure 1**), indicating that most of the detected Myxococcota are placed in the uncultured clades. The recovered Myxococcota bins showed larger genome sizes ranging from 6.6 to 10.7 Mb (average 9.2 ± 1.8 Mb) than general bacterial genomes such as *E. coli* (4.5–5.5 Mb), and similar genome sizes of myxobacteria were obtained compared to the previous studies (C. Wang et al., 2023). The 16S rRNA gene-based myxobacterial composition was similar to that obtained by metagenomic analysis (**Table S1 and Table S2**), indicating that the metagenomic bins of the predominant Myxococcota were correctly recovered from the activated sludge samples in this study.

**Figure 1.**
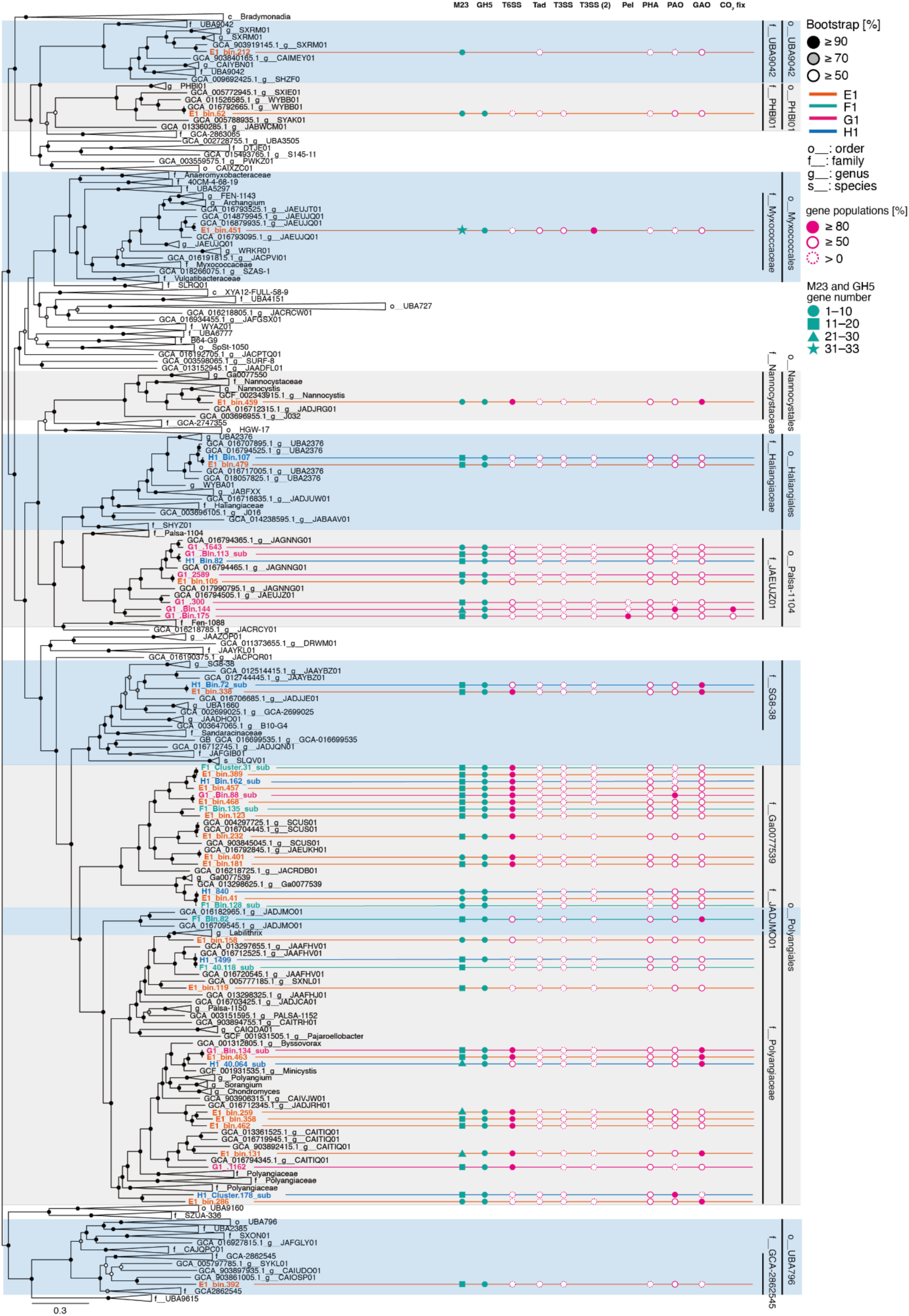
Genome tree of the phylum Myxococcota based on concatenated phylogenetic marker genes in GTDBtk 2.0.0 (ver. 207). The black, gray, and white circles indicate the ultrafast bootstrap (1000 replicates)-supported probabilities at >90%, >70%, and >50%, respectively. The magenta circles, solid lines with white circles, and dashed white circles show gene populations of ≥ 80%, ≥ 50%, and > 0%, respectively. Green circles, squares, triangle, and stars indicate detected numbers of M23 and GH5 families ranging from 1–10, 11–20, 21–30, and 31–33, respectively. M23 and GH5: M23 and GH5 families, T6SS: Type VI secretion systems, Tad: tight adherence (Tad)-like systems, T3SS: Type VI secretion systems, T3SS(2): Type VI secretion systems (2), Pel: Pel genes, PHA: polyhydroxyalkanoate production, PAO and GAO: polyphosphate and glycogen accumulating organism-related metabolism, and CO_2_ fix: CO_2_ fixation pathways with ribulose 1,5-bisphosphate carboxylase/oxygenase (RuBisCO), Calvin–Benson–Bassham (CBB) cycle, carotenoid metabolism, and bacteriochlorophyll production.

**Table 1.**
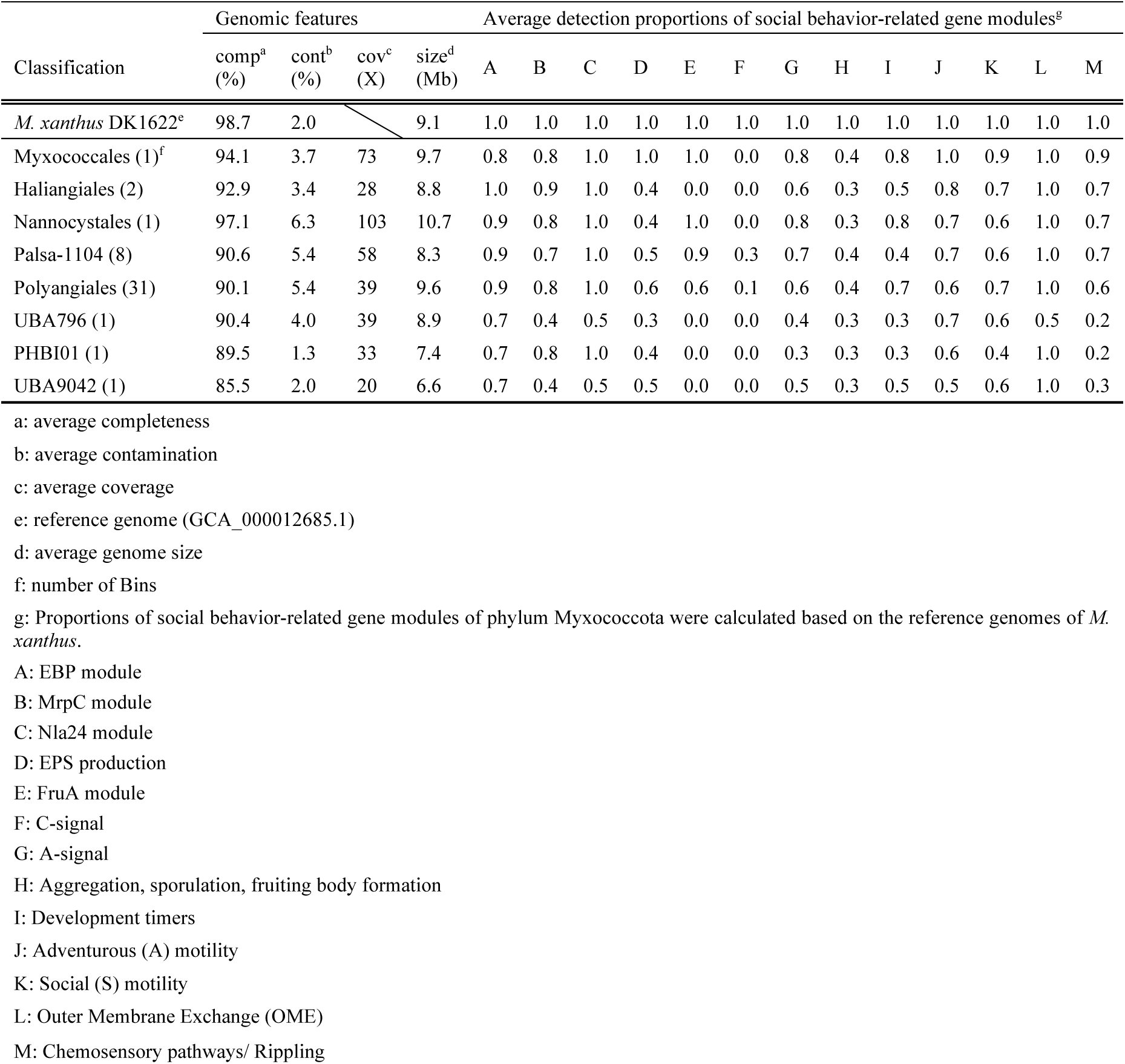
Identified genes relevant to social behaviors of the Myxococcota metagenome-assembled genomes obtained from activated sludge systems.

### 3.2 Predatory functions

Typical myxobacteria, such as *M. xanthus* can predate other bacteria (*e.g.*, *E. coli* and *Saccharomyces cerevisiae*) by producing degradative enzymes (Thiery et al., 2022). To evaluate their predatory potential, principal component analysis (PCA) was performed using the number of identified peptidases and glycoside hydrolases in the Myxococcota bins (**Table 1**, **Fig. 2, Table S3, and Table S4**). The results indicated that the order Polyangiales possesses many M23 family genes, including peptidase enzymes that cleave peptide bonds, such as the peptidoglycan of bacterial cell walls (Razew et al., 2022) (**Fig. 2A**). Furthermore, bacteriocins and autolysins which belong to the M23 family have effects such as inhibition of protein synthesis and antibacterial activity against other bacteria (Cleveland et al., 2001). Regarding glycoside hydrolases, many genes of the GH5 family, which contain cellulases and exoglucanases, were identified in the Palsa-1104 and Polyangiales bins (**Fig. 1 and Fig. 2B**). Our previous study revealed that Myxococcota bacteria are predominant in activated sludge systems in which wastewater treatment was stable (Kuroda et al., 2023). In addition, Myxococcota have been reported to be active predators in activated sludge processes (Zhang et al., 2023). These results imply that Myxococcota can prey on several operational problem-causing bacteria, such as bulking-causing filamentous bacteria, to maintain a stable environment for wastewater treatment. Some Myxobacteria are reported to be capable of producing *β*–barrel-glucanase (GluM), which allows them to prey on fungi with β-1,6-glucane in the cell wall (Li et al., 2019). Therefore, the myxobacteria belonging to Palsa-1104 and Polyangiales obtained in this study may be able to prey on fungi in addition to bacteria in activated sludge.

**Figure 2.**
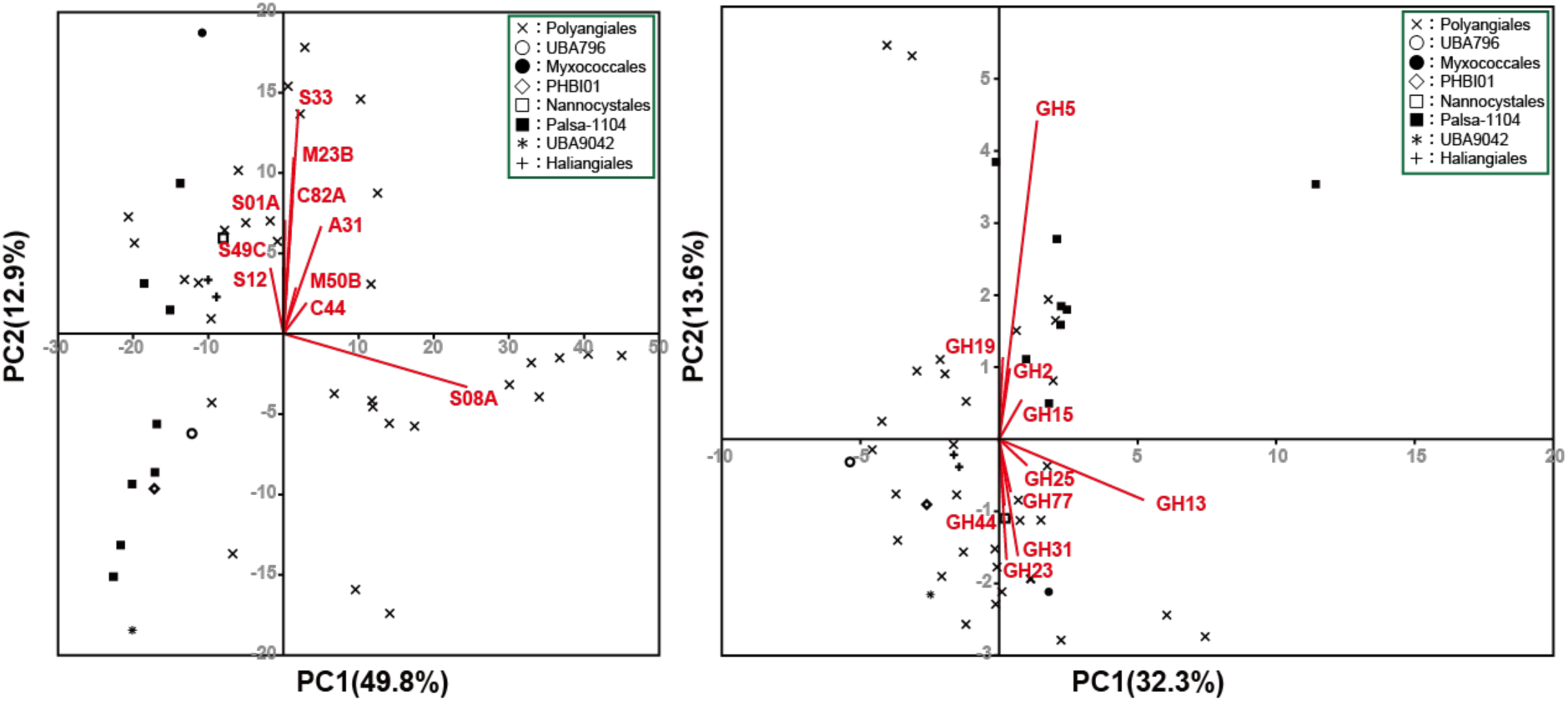
Principal component analysis (PCA) of numbers of genes of (A) peptidase and (B) glycoside hydrolase. Each number with alphabet indicates family of peptidase or glycoside hydrolase.

In addition, *M. xanthus* DK1622 possesses cell contact-dependent predatory functions that digest bacterial cells to obtain nutrients (Seef et al., 2021). The predatory functions require several secretion systems such as T2SS, T3SS, T4SS, and T6SS (Thiery et al., 2022) and it has been predicted that the *M. xanthus* DK1622 utilizes two types of T3SS and “Kill complex” of Tad-like secretion for the export of several digestive enzymes from the cells (Seef et al., 2021; Thiery et al., 2022). To estimate the functions of the bins of this study, amino acids-based homology search with the CDSs of *M. xanthus* DK1622 was performed with the threshold of 1e^-5^ ≥ e-value (**Fig. 1**, **Fig. 3, and Table S5**). The results clearly showed that ≥ 50% of bins in the family Myxococcaceae possess the homologs of Tad-like secretion, T3SS, and T3SS (2), which were consistent with our previous study (Kuroda et al., 2023). Furthermore, bins belonging to Nannocystis, Palsa-1104, Polyangiales, Ga0077539, SCUS01, Polyangiaceae, Minicystis, and SG8-3 encoded T6SS. Mutations in *M. xanthus* T6SS did not affect the cell contact-dependent predation of *E. coli* (Seef et al., 2021); therefore, these taxa may not have cell contact-dependent predatory functions or may have different predatory mechanisms from those of the family Myxoccocaceae.

**Figure 3.**
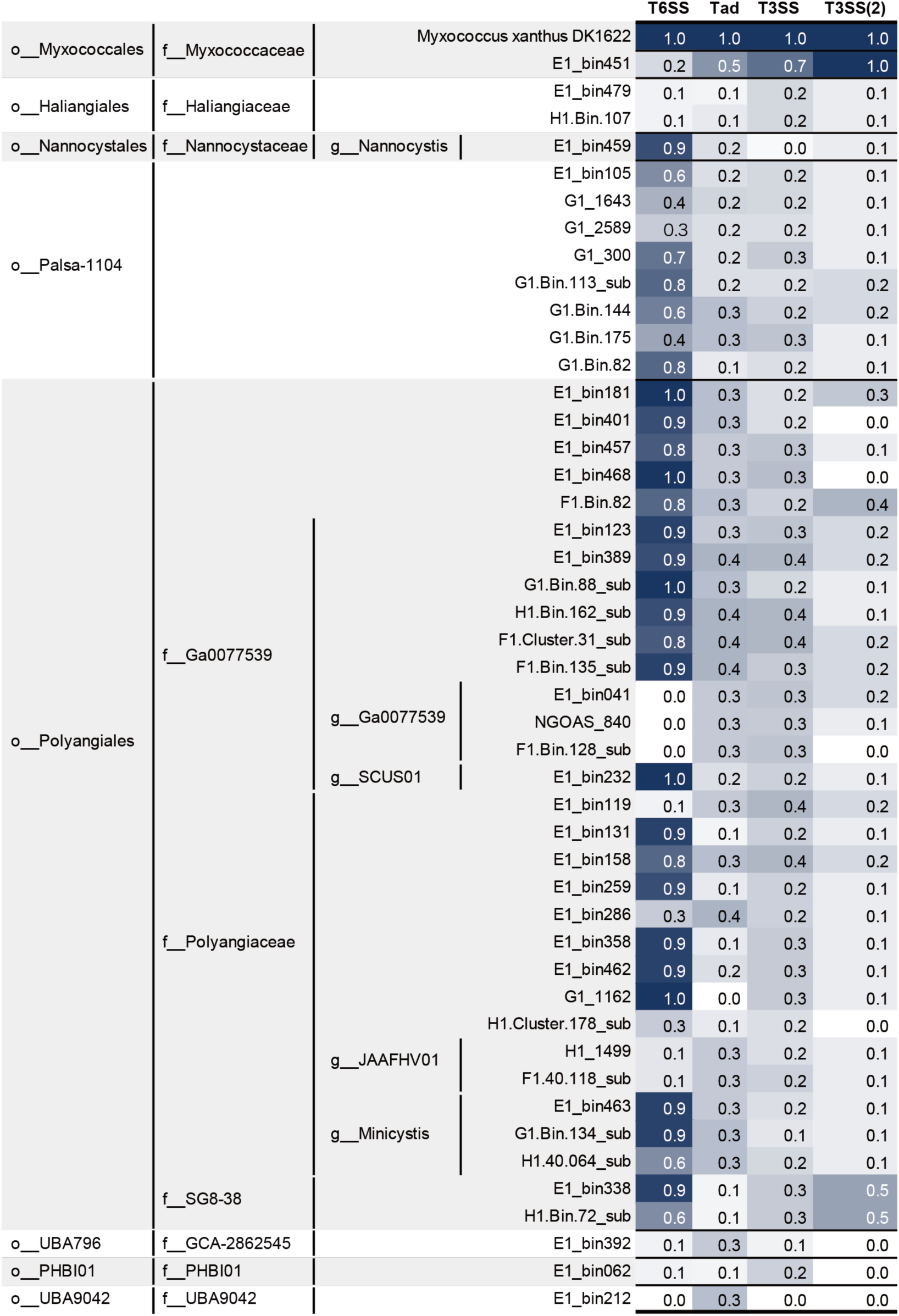
Possible cell contact-dependent predatory functions of the metagenomic bins of the phylum Myxococcota in activated sludge systems. The Type VI secretion systems (T6SS), tight adherence (Tad)-like systems, and Type III secretion systems (T3SS) were annotated with the genome of *Myxococcus xanthus* DK1622 (GCA_000012685.1) at the thresholds of ≥25% amino acid identity, ≤1e^-5^ e-value, and ≥50% query coverage per subject. The number of each secretion system indicates the gene proportions in the metagenomic bin. Gene populations were calculated based on the number of genes in each family. The detailed annotation results are listed in **Tables S5**.

### 3.3 Social behaviors

The social behavior-related genes of the phylum Myxococcota comprise 13 modules (enhancer-binding protein (EBP), MrpC, Nla24, EPS production, FruA, A-signal, C-signal, aggregation, sporulation, fruiting body formation, development timer, adventurous motility, social motility, outer membrane exchange, and chemosensory pathways/ribboning) (Murphy et al., 2021). In this study, the presence of gene clusters related to motility and fruiting body formation was confirmed by amino acid sequence-based homology searches using *M. xanthus* DK1622 as the reference genome (**Table 1, Table S2, and Table S6**). Myxococcota genomes in the activated sludge systems shared (> 0%) EBP, MrpC, Nla24, EPS production, A-signal, aggregation-sporulation-fruiting body formation, developmental timers, adventurous motility, social motility, outer membrane exchange, and chemosensory pathways/rippling modules. *M. xanthus* moves on solid surfaces using type IV pili, which have independent flagellar motility and frequently gets in contact with other myxobacterial cells (Chen and Nan, 2022). TraA and TraB on the outer membrane act as cell surface receptors that recognize whether a neighboring cell is clonal or not (Vassallo et al., 2015). The OME exchanges outer membrane proteins, lipids, and toxic compounds across the coupled outer membrane. When myxobacteria, which have different types of toxins depending on their phylogeny, exchange proteins with non-selfcells, each cell recognizes different myxobacteria through immunity proteins (Sah and Wall, 2020). The combination of TraA and OME recognition allows clonal cells to aggregate and form social communities. As homologs of TraA and TraB have been identified in Myxococcota genomes, they use a clonal cell-recognition mechanism similar to that of *M. xanthus*. Pili are important proteins that regulate the S motility. Through annotation of pili genes relevant to S motility, only the homolog of pilA was found in bins belonging to Palsa-1104 and Polyangiales. PilA synthesizes pili and drive cells in a forward-pulling motion. However, in other Myxobacteria such as *M. xanthus* (Sharma et al., 2018), pilB and pilT have been reported to perform the same functions as pilA. In addition, pilB and pilT have been identified in several bins of Myxococcota that have no pilA, suggesting that other genes, such as pilB and pilT, may complement the role of pilA in S motility. In contrast, FruA and C signals were only detected in bins belonging to Palsa-1104 and Polyangiales. The FruA module is relevant for controlling aggregation associated with fruiting body formation and sporulation (Campbell et al., 2015), and the C-signal module contains the PopC, PopD, and CsgA genes, which control the onset of fruiting body formation (Boynton and Shimkets, 2015). The CsgA gene is a precursor of the C-signal, which is necessary for fruiting body formation and sporulation, and promotes sporulation outside the fruiting body by excessive secretion. In addition, PopC protease hydrolyzes CsgA to produce C-signals. PopD inhibits PopC protease activity by forming a complex with PopC C-signal, which recognizes neighboring myxobacterial cells and activates signaling (ActA, ActB, and FruA) after secretion outside the outer membrane. Therefore, Palsa-1104 and Polyangiales may not have been able to form fruiting bodies because of the lack of CsgA genes in this study. To the best of our knowledge, no reports are available on fruiting body formation in the orders Palsa-1104 and Polyangiales detected in activated sludge, suggesting that these taxa do not require fruiting body formation in their habitats. Furthermore, we could not identify the complete set of genes relevant to fruiting body formation from the 46 activated sludge Myxococcota bins, including the family Myxococcaceae (**Table 1, Table S2, and Table S6**). This may be explained by the presence of rich nutrients in wastewater; therefore, fruiting body formation is not required for adaptation to activated sludge environments.

### 3.4 Metabolic potentials

To assess the metabolic information of the phylum Myxococcota in activated sludge environments, carbon, nitrogen, and sulfur metabolisms were annotated using the GhostKOALA annotation tool (**Fig. 4A and Table S7**). Most Myxococcota bins had almost complete metabolic pathways for glycolysis and the TCA cycle. Bins belonging to Myxococcales, Haliangiales, Nannocystales, Palsa-1104, Polyangials, and UBA796 have encoded β-N-acetylhexosaminidase genomes, and the enzyme has been reported to be involved in chitin degradation and the formation of chitinase derivatives in marine chitin-degrading bacteria (Keyhani and Roseman, 1996). In addition, some bins in orders Polyangiales and Palsa-1104 possess the beta-glucosidase, which is relevant for cellulose degradation. *Bacteria* in the genus *Sorangium* belonging to the family Polyangiaceae, are known to be cellulose-utilizing myxobacteria (Schneiker et al., 2007). This suggests that the bins in G1_Bin.134_sub, E1_bin.463, and H1_40.064_sub may have cellulose availability because of their close phylogenetic placement with the genus *Sorangium*. In nitrogen metabolism, bins in Myxococcales, Haliangiales, Polyangiales, and Palsa-1104 have the genes nirK/nirS, NorB/C, and NosZ, suggesting that the Myxococcota bins could contribute to denitrification in activated sludge systems. As for sulfur utilization, genes involved in the sulfide oxidation and sulfur assimilation are shared in phylum Myxococcota except for the Myxococcales and PHBI01. Sulfide dioxygenases (SDOs), sulfide-quinone oxidoreductase (sqr), flavoprotein of sulfide dehydrogenase (fccB), and sulfite reductase (ferredoxin; sir) were annotated. Therefore, Myxococcota, except for Myxococcales and PHBI01, may catalyze the oxidation of S^0^ to sulfite, detoxify hydrogen sulfide, and reduce sulfite (Wu et al., 2017).

**Figure 4.**
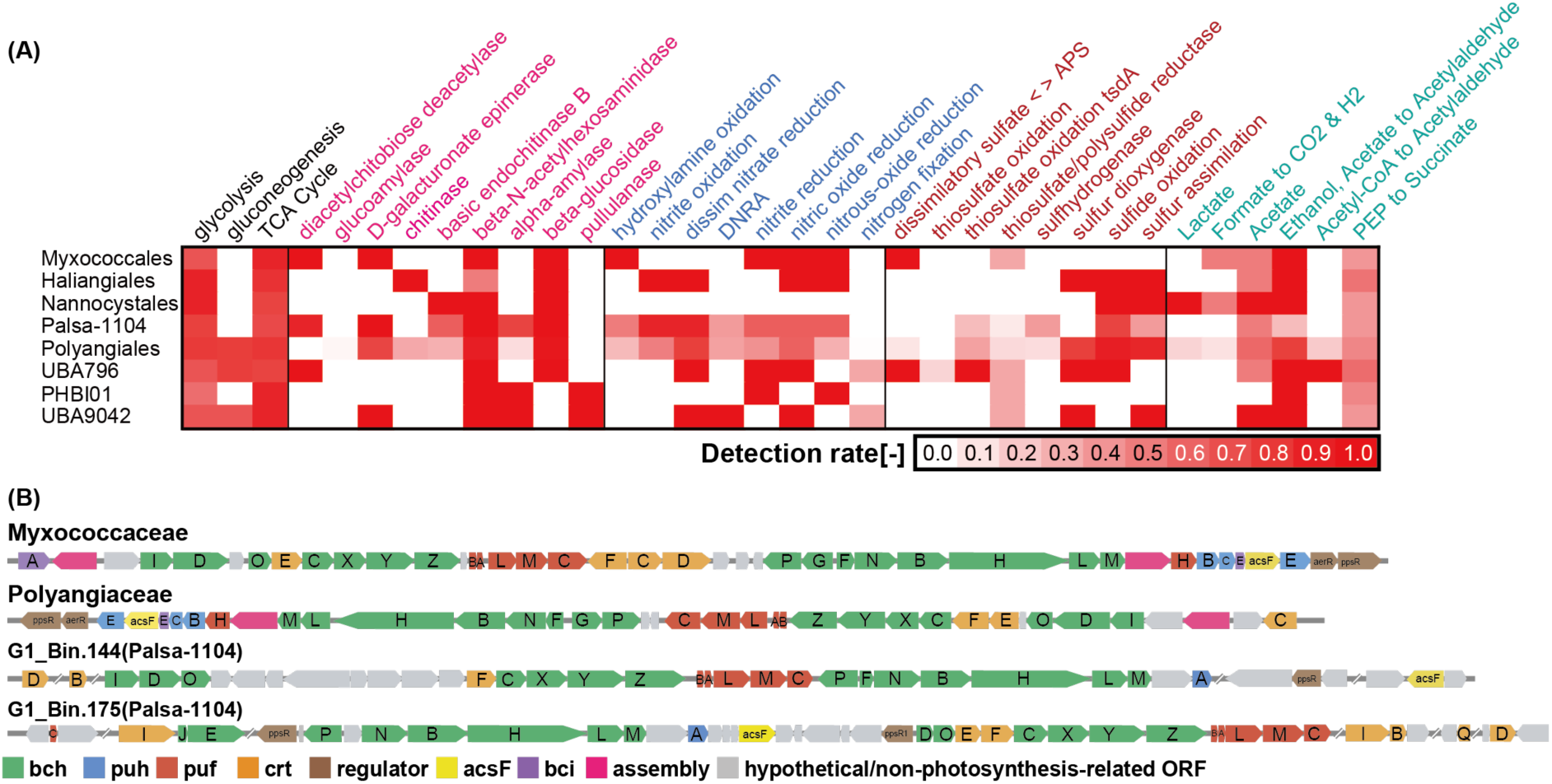
(A) Metabolic potentials of the metagenomic bins of the phylum Myxococcota in the activated sludge systems. (B) Potential phototrophic Myxocococcota (G1_Bin.144 and G1_Bin.175) in the activated sludge systems and photosynthetic gene clusters (PGCs) compared with known PGCs in a previous study (Huang et al., 2023). Genes of bacteriochlorophyll biosynthesis, reaction center proteins, reaction center assembly proteins, and carotenoid biosynthesis genes, are denoted as bch (green)/bci (purple), puf(red), puh (blue), and crt (orange), respectively. Hypothetical genes are in gray colour. Details of the annotation are provided in **Table S10**.

### 3.5 Biofilm formation

*M. xanthus* secretes extracellular polysaccharides during migration and colonization (Zhou and Nan, 2017). A recent study also identified pel genes in the MAG of Myxococcota in activated sludge systems (Dueholm et al., 2023). Pel is widely conserved in gram-positive and gram-negative bacteria, contributes to biofilm and pellicle formation, and mediates cell-cell interactions (Jennings et al., 2015). In this study, pel genes were identified in three bins (G1_300, G1_Bin.144, and G1_Bin.175) belonging to Palsa-1104. Conversely, no signal peptides were observed in the Pel genes in this study, suggesting that Pel genes are not secreted extracellularly into activated sludge and that Pel does not contribute to biofilm formation in activated sludge (**Fig. 1, Table S7, Table S8, and Table S9**). Genes relevant to polyhydroxyalkanoate (PHA) production are found in most Myxococcota genomes. PHA production has been reported to possibly promote biofilm formation in activated sludge (Chang et al., 2024). Therefore, Myxococcota in activated sludge may contribute to biofilm formation through PHA production.

In activated sludge systems, polyphosphate- and glycogen-accumulating organisms (PAOs and GAOs) are important bacteria that control biological phosphorus removal processes. These microorganisms form bacterial flocs during the phosphorous or glycogen. Controlling the abundance of PAOs is essential for effective removal of phosphorus from wastewater. PAO (such as *Ca*. Accumulibacter phosphatis) and GAO (such as *Ca*. Defluviicoccus tetraformis strain TFO71) have similar PHA metabolism (glgABCEX) and glycogen metabolism (phaABCEJ and fabDFG) genes (amino acid sequence similarity: up to 72%) when coexisting under anaerobic-aerobic conditions, which may be due to similar genetic adaptations as a result of selection pressure for survival in an enhanced biological phosphorus removal (EBPR) bioreactor (Nobu et al., 2014). Comparison of Myxococcota in this study with the metabolism of PAOs and GAOs showed no significant differences in the presence or absence of genes involved in PHA synthesis/degradation and glycogen metabolism (**Fig. 1 and Table S8**); therefore, physiological experiments of activated sludge Myxococcota are required to clarify their contribution to phosphorus removal during wastewater treatment. In previous studies, positive correlations were observed between mle1-27, other Myxococcota, and organic matter removal rates (Zhang et al., 2023). In addition, mle1-27 was positively correlated with the genus *Tetrasphaera* which is known as PAO. These results suggest that Myxococcota may be involved in the phosphorus removal during activated sludge processes.

### 3.6 Carbon fixation

In a recent study, metagenomic analysis revealed that the families Houyibacteriaceae, Myxococcaceae, Nannocystaceae, Polyangiaceae, Kuafubacteriaceae, and Sandaracinaceae in the phylum Myxococcota have metabolic pathways important for photosynthesis, including ribulose 1,5-bisphosphate carboxylase/oxygenase (RuBisCO), the Calvin-Benson-Bassham (CBB) cycle, carotenoid metabolism, and bacteriochlorophyll production (Li et al., 2023). *M. xanthus* is a heterotrophic bacterium with glycolytic and TCA cycles; however, this study has reported some Candidatus species. Kuafubacteriaceae and Myxococcaceae possess autotrophic metabolism with a near-complete CBB cycle (Li et al., 2023). In this study, G1_Bin.144 and G1_Bin.175, belonging to Palsa-1104 and recovered from G1, possessed ribulose RuBisCO and CBB cycles, suggesting that these bins might fix CO_2_ through the carbonate fixation pathway (**Fig. 1 and Table S10**).

A BLASTP homology search of the NCBI database identified the RuBisCOs in these Bins as type I RuBisCOs. Type I RuBisCO catalyzes carbon fixation reactions during photosynthesis and is widely conserved in plants and photosynthetic bacteria (Liu et al., 2023). In addition, biosynthetic pathways for carotenoids and chlorophyll were also identified, supporting the possibility of autotrophic metabolism in the Myxococcota bins (**Fig. 4B, Table S7, and Table S10**). Based on the annotations of the photosynthetic pathways, G1_Bin175 and G1_Bin144 had carotenoid and bacteriochlorophyll synthesis pathways. Carotenoids act as light-harvesting pigments that transfer energy to chlorophyll and are found in the vicinity of chlorophyll (Schoefs, 2002). In this study, G1_Bin.144 has several metabolic pathways capable of catalyzing the precursor bacteriochlorophylide and producing bacteriochlorophyll. G1_Bin.175 also lacks a part of the bacteriochlorophyll biosynthesis pathway; however, a gene that catalyzes bacteriochlorophyll production has been identified, suggesting that it may produce bacteriochlorophyll. These results suggest that some Palsa-1104 strains may have both heterotrophic and autotrophic metabolic pathways.

### 3.7 Prediction of secondary metabolism

A total of 533 BGCs were identified based on the evaluation of secondary metabolite biosynthesis genes using antiSMASH (**Table S11**). Nannocystales (26 BGCs per bin), UBA796 (19), Palsa-1104 (16), and Polyangiales (11) had higher numbers of BGCs per genome than the other orders of Myxococcota. In addition, BGCs classified as non-ribosomal peptide synthase (NRPS) and polyketide synthase (PKS) were mainly encoded by Palsa-1104 (50 BGCs) and Haliangiales (88 BGCs). Bacteriocins and aryl polyenes were frequently found in annotated BGCs. Bacteriocins are antibiotics that are conserved in a wide range of bacteria and act on closely related species (Cleveland et al., 2001). Myxobacteria exchange toxins as a recognition mechanism to avoid the grouping of different species during OME (Vassallo et al., 2015). Thus, bacteriocins are highly conserved among myxobacterial cells to identify themselves. Aryl polyenes, which are structurally similar to carotenoids, have also been reported to protect cells from damage caused by reactive oxygen species and promote biofilm formation by EPS in *E. coli*. EPS production by *M. xanthus* is important for smooth S cell movement and colony formation (Johnston et al., 2021; L. Wang et al., 2023). These results suggest that Myxococcota in activated sludge use bacteriocins during OME to avoid grouping with different species and promote EPS production. The fact that EPS production in *M. xanthus* is essential for aggregation, fruiting body formation, and gliding suggests that aryl polyenes may be involved in biofilm or polysaccharide production in Myxococcota. Secondary metabolic biosynthetic genes capable of producing aryl polyenes and bacteriocins were detected in some Polyangiales and Palsa-1104 strains in this study, suggesting that these substances are produced and used during OME and cell growth.

Furthermore, we identified the C domain in the NRPS and KS domain of PKS as the secondary metabolite biosynthetic genes of Myxococcota from activated sludge using the Natural Product Domain Seeker (NaPDoS) software (**Figure 5, Table S12, and Table S13**) (Ziemert et al., 2012). In the NRPS, 66 amino acid sequences annotated as bacitracin, bleomycin, cyclomarin, fengycin, gramicidin, iturin, microcystin, pksnrps2, pristinamycin, syringomycin, and tyrocidin were detected in the Myxococcota genomes. Fengycin and iturin may be useful biological pesticides in agricultural systems (Fira et al., 2018). In the PKS, we detected a total of 373 KS domain amino acid sequences; among these, 228 sequences were classified as fatty acid synthesis and polyunsaturated fatty acids. The remaining 145 amino acid sequences belong to virginiamycin, tylosin, tetronomycin, salinilactam, rifamycin, pyoluteorin, polyunsaturated fatty acids, nystatin, nidamycin, neocarzinostatin, mycocerosic acid synthase, megalomicin, lovastatin, and kirromycin. Virginiamycin and rifamycin, in the KS domain, have also been used as antibacterial agents (Batiha et al., 2020; Cocito, 1979). In contrast, the NRPS and PKS in this study were homologous to known amino acid sequences ranging from 21-47% (e-value: 7.0e-6–1.0e-100) and 23-73% (2.0e-6–1.0e-172), respectively, with most sequences having low homology. Therefore, the Myxococcota from the activated sludge in this study may probably have novel secondary metabolite biosynthesis genes. In addition, natural products derived from novel BGCs are considered promising therapeutics for animal and human health, and activated sludge Myxococcota can be considered potential novel resources.

**Figure 5.**
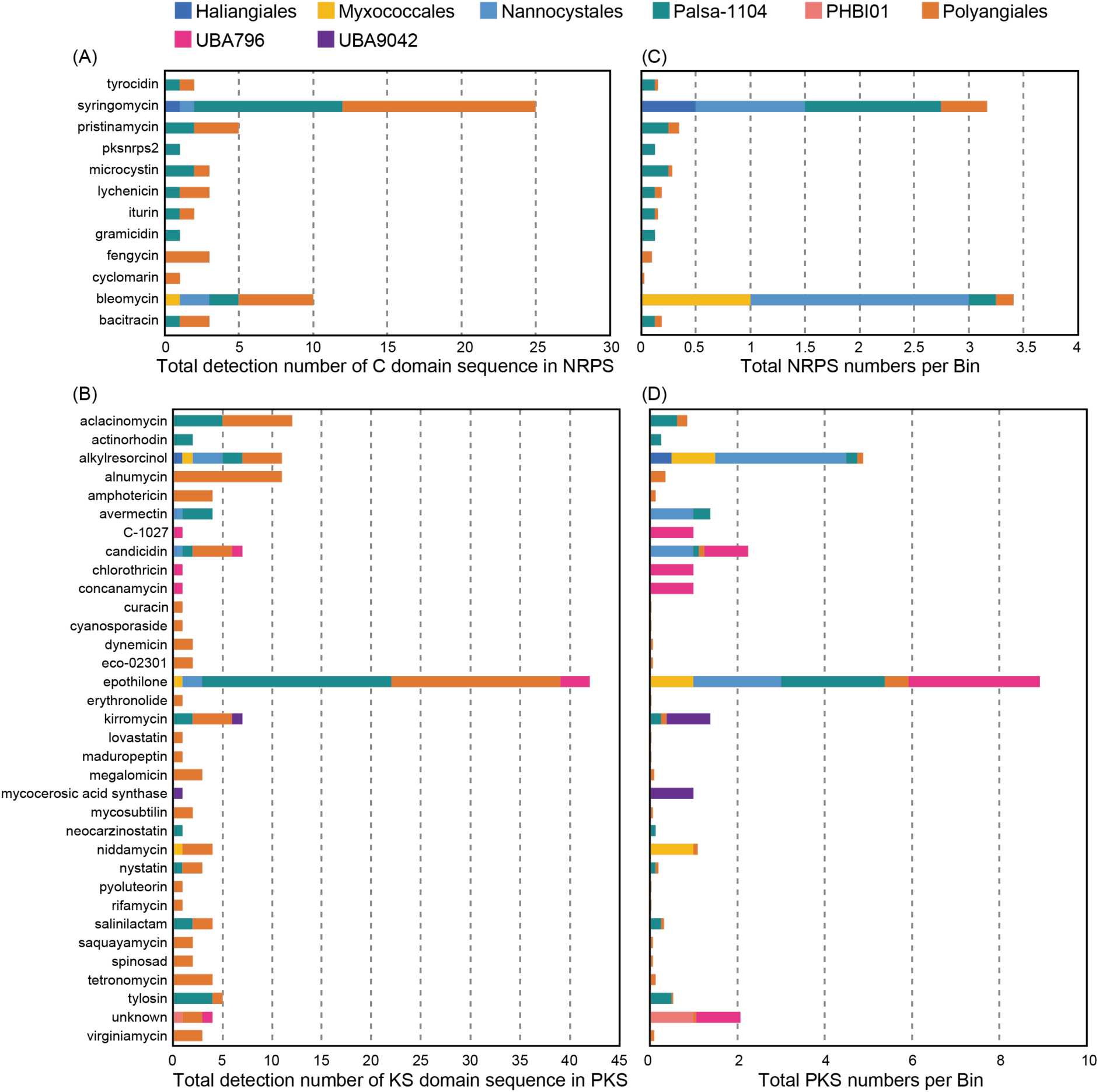
Results of identification of secondary metabolite genes (C-domain for non-ribosomal peptide synthase (NRPS) and KS-domain for polyketide synthase (PKS)) in Myxococcota metagenome-assembled genomes through NaPDoS software. (A) and (B) Total numbers of NRPS and PKS, respectively. (C) and (D) Numbers of NRPS per bin and PKS per bin, respectively.

## 4. Conclusions

To elucidate the ecophysiological characteristics of myxobacteria in activated sludge systems, we evaluated 46 metagenomic bins of the phylum Myxococcota recovered from four wastewater treatment plants. Through metagenomic analyses, we discovered novel functions of the activated sludge Myxococcota as follows: 1) Palsa-1104 and Polyangiales bins have the potential for cell wall degrading functions due to the presence of many of the family M23 and GH13 genes, 2) cell contact-dependent predatory functions have been conserved in the family Myxococcaceae, 3) fruiting body formation-related genes are lacking in the Myxococcota genomes, 4) two bins belonging to Palsa-1104 have phototrophic gene clusters with heterotopic metabolism, and 5) several novel secondary metabolite biosynthesis gene clusters are present in the Myxococcota genomes. These results indicate that the diverse ecophysiological functions of Myxococcota are hidden in activated sludge systems.

## Acknowledgments

This work is partly supported by the Cabinet Office, Government of Japan, Cross-ministerial Strategic Innovation Promotion Program (SIP), “Technologies for creating next-generation agriculture, forestry and fisheries” (funding agency: Bio-oriented Technology Research Advancement Institution, NARO). In addition, we gratefully acknowledge the collaborative works of members of the research consortium “Development of Novel Technology for Bioprocess Optimization in Smart Bio-Industry” of the SIP. Sequencing data submission was supported by the D-way Submission Portal provided by the DNA Data Bank of Japan (DDBJ). We thank Dr. Tomoyuki Hori, Dr. Tomo Aoyagi, Dr. Yuya Sato, Dr. Tomohiro Inaba, Dr. Hiroshi Habe, Dr. Hideyuki Tamaki, Dr. Yoshihisa Hagihara, Dr. Tomohiro Tamura, Ms. Chisato Shiiba, Ms. Maki Yanagisawa, Ms. Satomi Mori, Ms. Masayo Ogiso, Ms. Yumiko Kayashima, Mr. Toru Ishizuka, and Dr. Ling Leng (National Institute of Advanced Industrial Science and Technology (AIST), Japan) for their assistance. Hazuki Kurashita was supported by a Grant-in-Aid for JSPS Fellows (JP22KJ1452).

## Legends of supplementary tables

Table S1. Abundance of 16S rRNA gene sequences at phylum and order levels in the activated sludge systems from four wastewater treatment plants.

Table S2. Summary of the Myxococcota metagenomic bins obtained in this study.

Table S3. Number of glycoside hydrolase related genes from Myxococcota in activated sludge.

Table S4. Number of peptidase related genes from Myxococcota in activated sludge.

Table S5. Locus tags for possible cell contact-dependent predatory functions of Myxococcota in this study.

Table S6. Locus tags for genes encoding social behavior of the myxobacteria in this study.

Table S7. Summary of the metabolic functions of Myxococcota in this study.

Table S8. Locus tags for genes encoding PHA synthesis/degradation, polyphosphate/glycogen accumulation, and Pel synthesis in this study.

Table S9. Presence (1)/absence (0) of genes encoding PHA synthesis/degradation, polyphosphate/glycogen accmulation, and Pel synthesis in this study.

Table S10. Locus tags for genes relevant to photosynthetic gene clusters in this study.

Table S11. Summary of identified secondary metabolite biosynthesis genes using antiSMASH software.

Table S12. Summary of the identified C domain in non-ribosomal peptide synthase based on NaPDoS software.

Table S13. Summary of the identified KS domain in Polyketide synthase based on NaPDoS software.

